# RootQuant: Automated Root Trait Quantification from Minirhizotron Images Using Deep Learning

**DOI:** 10.64898/2026.07.07.737053

**Authors:** Kinjalk Parth, Sebastian Varela, Zhiru Liu, K. Michael Martini, Ashish Rajurkar, Dylan Allen, Scott McCoy, Jeremy Ruhter, Samuel Walker, Nigel Goldenfeld, Andrew D.B. Leakey

**Affiliations:** Institute for Genomic Biology, University of Illinois at Urbana-Champaign, Urbana, IL 61801, USA; Center for Advanced Bioenergy and Bioproducts Innovation, Urbana, IL 61801, USA; Center for Digital Agriculture, University of Illinois at Urbana-Champaign, Urbana, IL 61801, USA; Independent Researcher, Canelones 15800, Uruguay; Department of Physics, University of Illinois at Urbana-Champaign, Urbana, IL 61801, USA; Department of Plant Biology, University of Illinois at Urbana-Champaign, Urbana, IL 61801, USA; Institute for Sustainability, Energy and Environment, University of Illinois at Urbana-Champaign, Urbana, IL 61801, USA; Department of Crop Sciences, University of Illinois at Urbana-Champaign, Urbana, IL 61801, USA

**Keywords:** deep learning, root phenotyping, minirhizotron, maize, soybean

## Abstract

Quantifying root traits such as root length (RL) and root surface area (RSA) from minirhizotron imagery is a valuable approach for overcoming the phenotyping bottleneck that limits understanding and improvement of crop productivity, resource use efficiency and resilience in field experiments. However, current approaches remain labor-intensive, and deep learning (DL) methods suffer from limited generalization ability. We present RootQuant, an end-to-end DL model that simultaneously predicts RL and RSA directly from minirhizotron images using only whole-image trait values as supervision, thereby eliminating the need for pixel-level annotations. The model’s generalization ability was evaluated across species and fine-tuning configurations. The practical applicability of the model was further assessed under field conditions by converting image-derived RL estimates into volumetric root length density (vRLD). Using 118,191 maize and soybean images collected between 2009 and 2020, RootQuant trained on both species achieved an R^2^ of 0.90 and an RMSE of 2.9 mm for RL, and an R^2^ of 0.88 and an RMSE of 4.2 mm^2^ for RSA. The same mixed-species model generalized strongly across species, yielding an 8% relative improvement in R^2^ and a 30% lower RMSE on maize compared with the same architecture trained on a single species and applied zero-shot. Image-derived RL predictions converted to vRLD showed the expected depth-dependent decline in vRLD, as was also found by coincident destructive quantification of roots washed out of soil cores. By providing a generalist backbone model trained on a large dataset from two major crop species, RootQuant enables high-throughput simultaneous estimation of two relevant root traits directly from raw imagery without task-specific fine-tuning, thereby accelerating in situ root system analysis and phenotyping applications.

## 1 Introduction

Plant root systems mediate critical soil–plant interactions, including water and nutrient uptake, carbon allocation, and mechanical anchorage, and are therefore important to plant environmental adaptation and agricultural productivity [1], [2], [3]. Among root architectural traits, RL and RSA are particularly informative because they capture complementary aspects of soil exploration and rhizosphere processes [4]. RL influences the spatial extent of root foraging, whereas RSA determines the interface for water and nutrient exchange and microbial activity. Given this functional significance, crop improvement strategies have been proposed that target RL and RSA to improve drought resilience and nitrogen-use efficiency in crops [3], [5].

Minirhizotron imaging is the most widely used non-destructive method for studying root dynamics in field conditions [6], [7], [8]. Field campaigns involve the installation of tens to thousands of transparent access tubes into the soil profile. This allows specialized cameras to be inserted to capture 1,000s to 100,000s of images through the tube walls of the root system where it contacts the tube along the soil profile [9]. Recent advances in the number of access tubes installed and images captured further exacerbate the need for tools supporting automated analysis of minirhizotron imagery.

Traditional workflows rely heavily on manual or semi-manual tracing in proprietary software such as WinRHIZO Tron. These procedures are labor-intensive, time-consuming, and susceptible to operator subjective judgement, limiting reproducibility across large field studies [10]. In addition, the resulting annotations are stored in proprietary formats that preserve only accessible aggregated numerical traits (e.g., total root length, surface area per image), while the underlying segmentations, skeletons, and pixel-level annotations remain inaccessible. As a result, decades of historical minirhizotron datasets cannot be easily exploited by most modern DL approaches, which typically require dense pixel-level supervision for semantic learning.

Classical image-analysis pipelines based on color thresholding, filtering, and skeleton tracing further suffer from limited robustness under heterogeneous field conditions [11], [12]. Variations in soil texture, illumination, moisture, and image noise frequently degrade segmentation quality and introduce systematic bias into downstream trait estimation. These limitations constrain scalability and reduce the reliability of root phenotyping at field scale.

DL has transformed image-based plant phenotyping in recent years [13], [14], [15], [16] providing substantial improvements in robustness and automation compared with handcrafted computer-vision pipelines [17], [18]. However, most DL approaches for root imagery remain segmentation-driven and therefore depend on extensive pixel-level annotation [19], [20], [21], [22]. Reported work has demonstrated strong segmentation accuracy [19], [20], [21], [23], [24], but its applicability is restricted by the high cost of annotation and by limited access to compatible training labels in historical datasets. Moreover, these studies have been developed and evaluated on relatively small datasets [20], [23], [25] and single-species collections [21], [22], [23], limiting their ability to generalize across crop species, soil conditions, and field imaging campaigns.

The feasibility of predicting root traits directly from whole minirhizotron images has been demonstrated in [26], [27]. While these studies represent an important step toward annotation-efficient root phenotyping, they were conducted on relatively small datasets and focused primarily on RL estimation. Developing a scalable and generalizable framework capable of jointly predicting multiple traits across species remains a challenge. To address these challenges, we propose RootQuant, an end-to-end deep learning framework for the simultaneous prediction of RL and RSA directly from raw minirhizotron images using only whole-image trait values as supervision. In contrast to segmentation-based pipelines [19], [20], [21], [22], [23], [24], the proposed model framework does not require pixel-level masks, skeleton annotations, or root detection labels, enabling the utilization of large historical WinRHIZO Tron image archives without costly re-annotation. Furthermore, the framework incorporates mixed-species (MS) training, providing a unified approach that alleviates the need for crop-specific model fine-tuning.

This work is organized as follows: 1) A large minirhizotron dataset used for model development is described. 2) An end-to-end multi-output framework for jointly predicting RL and RSA from minirhizotron imagery is introduced, using whole-image trait values as supervision and eliminating the need for pixel-level annotations. 3) The generalization capability of the proposed framework is evaluated under different species-training modalities. 4) The model’s internal representation is analyzed to assess whether meaningful biological features are learned from images. 5) Finally, the biological relevance of the proposed approach is assessed by comparing field-scale volumetric root length density (vRLD) estimates derived from RootQuant predictions with independent soil-core measurements.

## 2 Materials and Methods

### 2.1 Image data acquisition and description

All minirhizotron images were acquired using Bartz BTC-100X minirhizotron imager systems inserted into transparent acrylic tubes installed in field plots at the research farms of the University of Illinois Urbana-Champaign, which is in the primary Midwest U.S. growing region of maize and soybean; a detailed description of the process can be found in [9]. The imaging system produces RGB images. Maize images were acquired at a resolution of 640 × 480 pixels with a fixed field of view of 1.8 cm × 1.3 cm per frame [9], while soybean images were acquired at a resolution of 754 × 510 pixels [28], [29]. Considering the optical depth-of-field of approximately 2 mm of soil, each image from maize represents an estimated soil volume of 0.468 cm^3^ (1.8 × 1.3 × 0.2 cm). This imaging volume was later used for the conversion of image-level predictions into vRLD estimates, as is common practice in minirhizotron [29], [30].

The complete dataset comprised 118,191 annotated minirhizotron images collected between 2009 and 2020 from soybean and maize field experiments (Fig. 1). The large temporal and cross-species variability represented in the dataset enabled evaluation of both single-species and MS model generalization. Soybean datasets contributed 99,934 images acquired during 2009–2011, while maize datasets contributed 18,257 images acquired during 2016, 2019, and 2020. The dataset included substantial variability in species, seasons, soil conditions, and root densities, thereby providing a suitable benchmark for evaluating both the single-species and MS model generalization ability.

**Figure 1:**
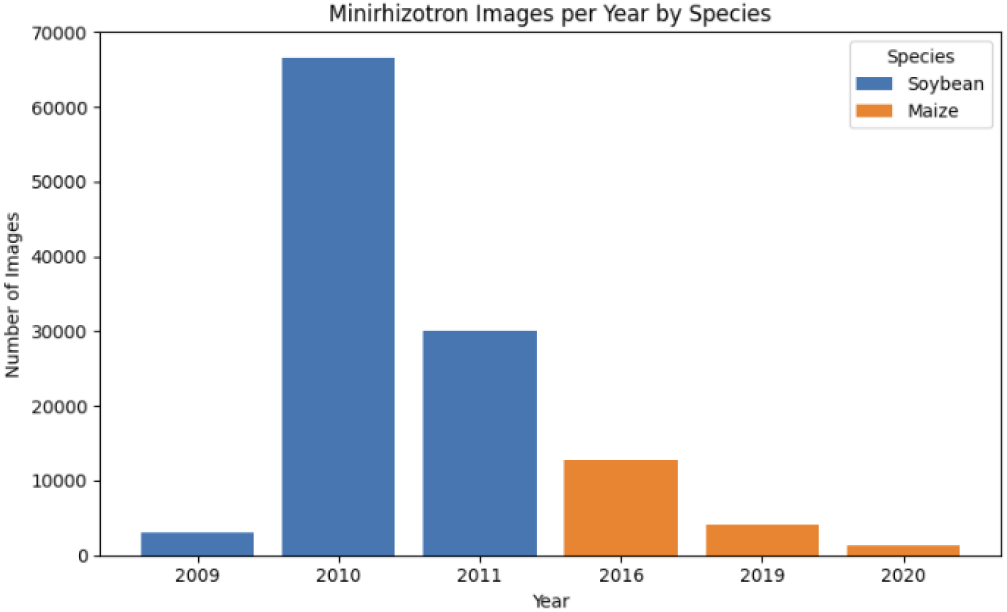
Distribution of minirhizotron images used for model development across years and crop species. The total number of images collected per year for soybean (2009–2011) (blue) and maize (2016, 2019, and 2020) (orange).

Each image was associated with two continuous whole-image target variables: RL (mm) and RSA (mm^2^). These labels were generated by trained human operators who manually traced visible roots within each image using the WinRHIZO Tron [31]. However, because WinRHIZO Tron stores annotations in proprietary formats, only the final numerical trait values per image were accessible.

A substantial proportion of the dataset consisted of zero-root images, in which no visible roots were present and both RL and RSA values were exactly zero. These samples were intentionally retained because they reflect the natural spatial and temporal distribution of roots observed in field minirhizotron campaigns. Excluding zero-root images would artificially bias the data distribution toward root-positive samples and would prevent the model from learning biologically realistic absence conditions.

### 2.2 Model architecture

RootQuant is an end-to-end supervised deep learning framework designed to predict RL and RSA directly from minirhizotron imagery. The model receives a single RGB minirhizotron image as input and simultaneously outputs two continuous values corresponding to RL (mm) and RSA (mm^2^) (Fig. 2). Importantly, training relies exclusively on whole-image numerical trait values exported from WinRHIZO Tron, without requiring segmentation masks, root skeletons, or keypoint annotations.

**Figure 2:**
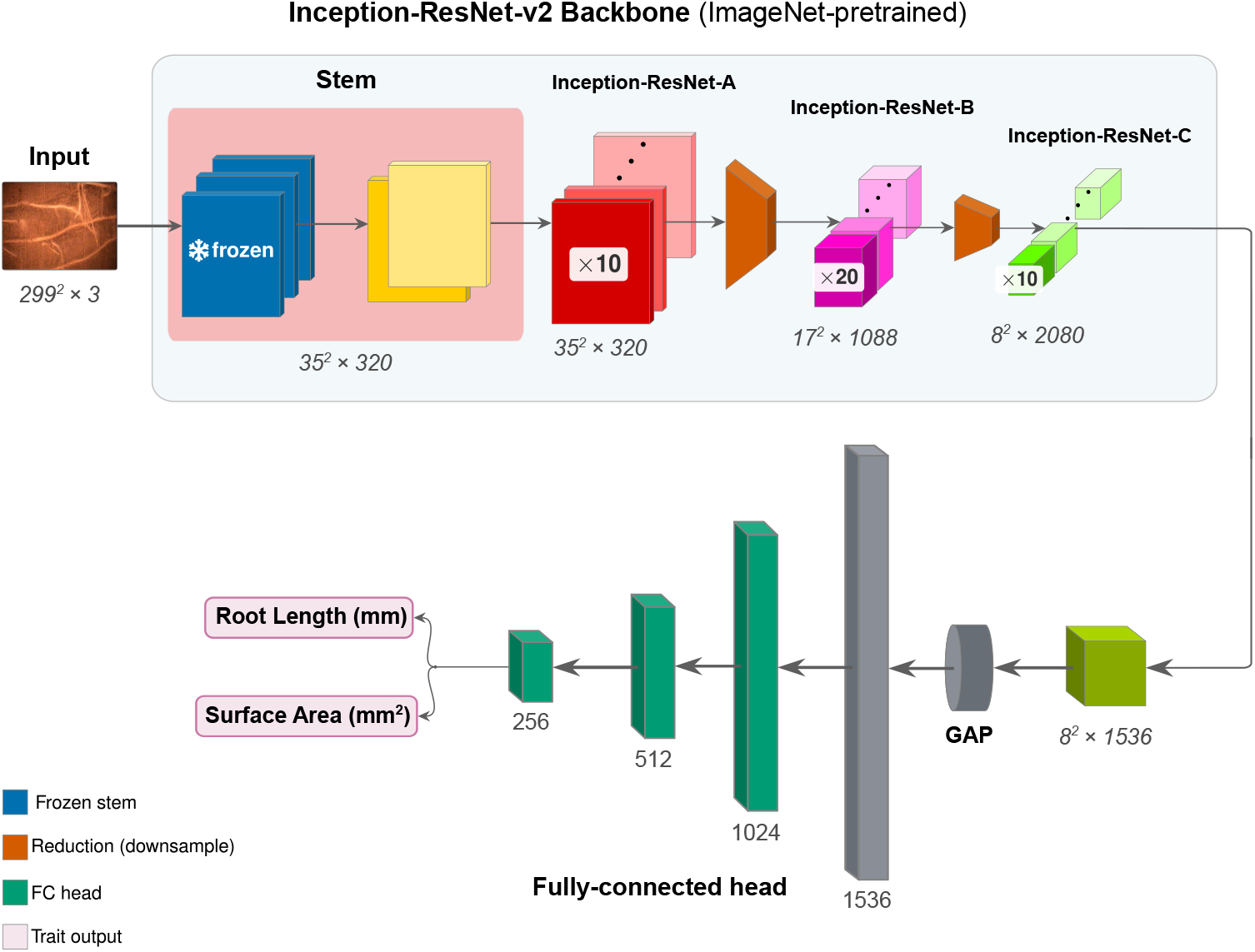
Overview of RootQuant architecture. Minirhizotron RGB images were resized to 299 × 299 pixels and processed using an ImageNet-pretrained Inception-ResNet-V2 backbone. Early stem layers (blue) were frozen to preserve transferable low-level image features, while deeper Inception A, B, C and reduction blocks (orange) were fine-tuned for root feature extraction. The resulting feature representation was passed to a custom dual-output regression head (dark green) that simultaneously predicted RL and RSA in z-score-normalized space.

RootQuant uses the ImageNet-pretrained Inception-ResNet-V2 [32], [33] as the visual feature extractor. This backbone combines residual connections [34] with multi-scale Inception modules [33], enabling efficient extraction of hierarchical image features across multiple receptive fields. Preliminary experiments showed that Inception-ResNet-V2 provided stronger feature representations than alternative architectures for heterogeneous minirhizotron imagery. Images originally captured with the Bartz BTC-100X minirhizotron imager system were resized to 299 × 299 pixels to match the network input requirements.

To preserve transferable low-level image features, the earliest convolutional stem layers of the backbone were kept frozen during all training configurations. All remaining convolutional blocks were used as trainable feature extractors, producing a 1536-dimensional representation for each image after global average pooling.

The extracted image features were passed to a custom fully connected regression head designed for simultaneous prediction of RL and RSA within a multi-task learning framework. The regression head consisted of multiple fully connected layers combined with Batch Normalization, ReLU activations, and dropout regularization.

Progressively decreasing dropout rates were applied across layers to reduce overfitting while preserving task-specific feature learning in deeper stages of the network. The final output layer used linear activation functions, allowing unconstrained continuous predictions that were later transformed back into physical units during inference.

### 2.3 Data preprocessing and model implementation

Root trait distributions were strongly right skewed, with most images containing either no visible roots or small root systems and a minority containing dense root structures. To stabilize optimization and prevent the loss from being dominated by large-value samples, target normalization was applied before training. The primary configuration used z-score normalization independently for RL and RSA based on statistics computed from the training split. Normalization statistics were computed exclusively on training data and stored with the trained model checkpoints to ensure consistent inverse transformation during validation, testing, and inference. The dataset was divided into training, validation, and test subsets using stratified random sampling based on root length distribution, with zero-root images treated as a separate stratum. This ensured preservation of the strong zero inflation and overall trait distribution across all partitions.

The final split contained 89,186 training images, 11,445 validation images, and 17,560 test images. Approximately 72–74% of images across all partitions contained no visible roots, reflecting the natural distribution observed in field minirhizotron campaigns.

Only light image augmentation was applied during training to preserve biologically relevant root textures and avoid unrealistic transformations of soil-background structure. Full details on image augmentation are available in supplementary information.

Model training used a multi-task objective [35] based on the weighted sum of two Huber losses [36], one for RL and one for RSA prediction. Equal weighting was assigned to both traits because normalization placed them on comparable numerical scales. Huber loss was chosen since it provides robustness to outliers and noisy labels while retaining sensitivity to small prediction errors. This was particularly important given the long-term, multi-operator nature of the dataset, which introduced variability associated with manual tracing, software-version differences, and occasional unit-conversion inconsistencies in historical WinRHIZO Tron exports.

Training was implemented in PyTorch [37] using the AdamW optimizer with OneCycle learning-rate scheduling. Mixed-precision BF16 training, gradient clipping, dropout regularization, and Batch Normalization were used to improve optimization stability and computational efficiency. All experiments were trained on a single NVIDIA A100 GPU.

Different model configurations were evaluated to characterize model generalization across species. The primary configuration, referred to as the MS generalist, was trained jointly on combined soybean and maize datasets. This configuration evaluated whether a single model could generalize across species without requiring species-specific fine-tuning. Starting from Inception-ResNet-V2 weights with frozen stem layers, all remaining backbone layers and the regression head were trained jointly using the combined soybean and maize training dataset (89,186 images). Training used z-score target normalization computed from the combined training split, and minibatches were generated through uniform random sampling without species balancing or oversampling. Model selection was based on the highest combined R^2^ obtained on the MS validation set, and final evaluation was performed on the held-out MS test dataset.

Additional evaluations investigated the cross-species transferability of a soybean model to maize. First, a soybean-only model was trained using the same pipeline and strategy but restricted to soybean training images. This model was then directly evaluated on maize images without additional training, constituting a zero-shot transfer (ZST) experiment and providing a lower-bound estimate of cross-species generalization ability.

Two fine-tuning strategies were subsequently evaluated to quantify the extent of adaptation required for transferring soybean-learned representations to maize imagery. In the fully connected fine-tuning (FCFT) configuration, the pretrained soybean backbone remained frozen and only the regression head was fine-tuned using maize training images. This experiment assessed whether species differences could be accommodated through recalibration of the output layers alone. In the full fine-tuning (FFT) configuration, all backbone layers except the frozen stem were fine-tuned using maize data, allowing broader feature adaptation across species. To reduce catastrophic forgetting of soybean-learned representations, backbone layers were optimized using a learning rate ten times smaller than that applied to the regression head.

All evaluations were performed on held-out test datasets, and all metrics were computed in the original physical units after inverse transformation of normalized predictions. RL was evaluated in millimeters (mm) and RSA in square millimeters (mm^2^). Training performance was monitored using the validation dataset after every epoch. The primary model-selection criterion was the combined coefficient of determination (2). The checkpoint with the highest validation combined *R*^2^ was retained, and early stopping was applied when validation performance failed to improve for 30 consecutive epochs. Final evaluation was performed on the held-out test set using *R*^2^ (1), RMSE (3), and MAE (4) computed in original physical units after inverse target normalization.

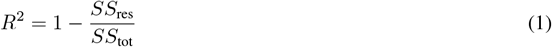

where *R*^2^ is the coefficient of determination.

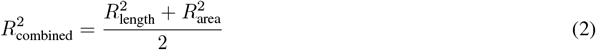

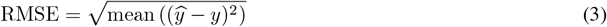

where RMSE is root mean squared error.

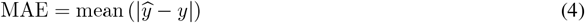

where MAE is mean absolute error.

### 2.4 Model’s internal representation

Whole-image regression models introduce a fundamental interpretability challenge – the model is optimized to minimize error on global scalar targets without any explicit spatial supervision. Because training is driven solely by image-level RL and RSA values derived from WinRHIZO Tron, there is no constraint forcing the network to optimize internally to root structures. This creates a setting in which the model may achieve high predictive accuracy by exploiting spurious correlations rather than true signal. In particular, non-root structures and conditions such as soil texture variation, illumination gradients, or systematic acquisition differences can become predictive shortcuts. This is consistent with the well-established phenomenon of shortcut learning in deep neural networks, where models preferentially rely on easy-to-learn but semantically irrelevant features when they are statistically predictive [38].

To assess whether the learned representations are grounded in root phenotypes or driven by such shortcuts, a combined attribution framework based on Grad-CAM [39] and LayerCAM [40] is implemented. Grad-CAM provides coarse, semantically meaningful localization maps by propagating gradients from the output back into the final convolutional layers, thereby highlighting regions that most strongly influence the prediction. These maps are primarily useful for qualitative inspection, offering an interpretable visualization of where the model appears to focus in the image. In contrast, LayerCAM extends this idea by computing activation-weighted gradient responses across convolutional feature maps, producing higher-resolution, pixel-level attribution maps. This finer spatial fidelity enables not only visualization but also quantitative measurement of how much predictive evidence is concentrated in biologically meaningful regions.

The evaluation utilizes 726 ground-truth binary root masks, where binary annotations define the spatial extent of root versus background pixels and therefore the biologically relevant regions of interest. Attribution maps are then generated using both methods: Grad-CAM is applied to the final convolutional block to produce interpretable overlays suitable for visual validation, while LayerCAM is applied across intermediate feature maps to obtain detailed spatial attribution distributions. These LayerCAM maps are used for quantitative analysis by computing the total attribution energy falling within root regions, defined as the sum of attribution values over all pixels inside the root mask. This can be expressed as *E*_*root*_ = ∑ _(*i,j*)ϵ*root*_ *A*_*i,j*_, where *A*_*i,j*_ denotes the attribution value at pixel (*i, j*). This value is normalized by the total attribution energy in the image to obtain a proportional measure of how much model evidence is concentrated on root structures.

To determine whether this concentration is meaningful, the in-mask attribution is compared against multiple baselines, including random area-matched masks that preserve the same coverage fraction but are spatially uninformative, as well as shape-preserving spatial shift controls that maintain mask geometry while displacing it away from true root locations. These comparisons allow separation of true localization signal from trivial effects of mask size or structure. In addition to energy-based metrics, a pointing game evaluation is performed by identifying the peak activation location in the LayerCAM map and testing whether this location falls within the ground-truth root mask. Given that root structures are sparse, the random baseline for the pointing game is the coverage fraction of the tolerance-banded root mask (∼19% of the frame with 5 px padding, ∼9% un-padded); a moderate pointing score is therefore expected for a correctly localized map.

Finally, all metrics are aggregated across the full evaluation dataset (i.e., 726 images) to obtain stable estimates of localization performance. This includes reporting mean enrichment ratios, distributions of per-image attribution concentration, and variability across samples. Together, these analyses provide a multi-level assessment of whether the model’s learned representations are genuinely aligned with root structures or instead reflect shortcut reliance on non-biological image structure.

### 2.5 Field-scale evaluation and validation of vRLD

To evaluate the agronomic relevance of RootQuant predictions, image-level RL estimates were converted into volumetric vRLD and compared against independent soil-core measurements obtained from maize field experiments (Fig. 3). Four minirhizotron tubes (two tubes in each of the two central rows) are located near the center of each 4-row plot. Minirhizotron images were acquired along obliquely installed access tubes using a BTC-100X scanner. Each image was assigned an along-tube position based on a layer index (L) with a fixed step (*δ* = 1.3462 cm). For each tube, the installation inclination (*ϕ*) and the distance from the tube opening to the viewing window were recorded. Vertical soil depth (z) was calculated from the along-tube position, the known hook spacing (1.27 cm), and the sine of the complement of *ϕ*, excluding images from above-ground sections or depths beyond 100 cm. Images were then binned into five 20 cm depth strata. RootQuant provided per-image predictions of root length (RL, mm). These were converted to volumetric root length density (vRLD, cm root cm^-3^ soil) by dividing the predicted length (converted to cm) by the imaged soil volume (width 1.8 cm × height 1.3 cm × depth 0.2 cm = 0.468 cm^3^). For comparison, independent soil cores (volume = *π* × (2.54 cm)^2^ × 20 cm = 405.37 cm^3^) were collected from four positions near the minirhizotrons within the central rows of each plot. For each sample, roots and soil were separated by washing, then root samples were taken to the lab and root length was determined by floating roots in water-filled transparent acrylic trays to minimize overlapping roots before scanning with a twin-sided flatbed scanner (Epson V800). The scanned image was analyzed using commercially available WinRHIZO software (Regent Instruments, Quebec, Canada). Finally, core vRLD was calculated as root length divided by core volume. Image-derived vRLD was averaged per (plot, tube, depth stratum) and then across tubes per (plot, depth). Core-derived vRLD was aggregated per (plot, depth), yielding 75 paired measurements (15 plots × 5 depth strata) for comparison. This field-scale comparison provided an independent validation of the biological consistency of RootQuant predictions beyond 2D image-level regression accuracy. A detailed description of the soil core and RootQuant vRLD calculation and the along-tube to vertical depth conversion is provided in Supplementary Materials.

**Figure 3:**
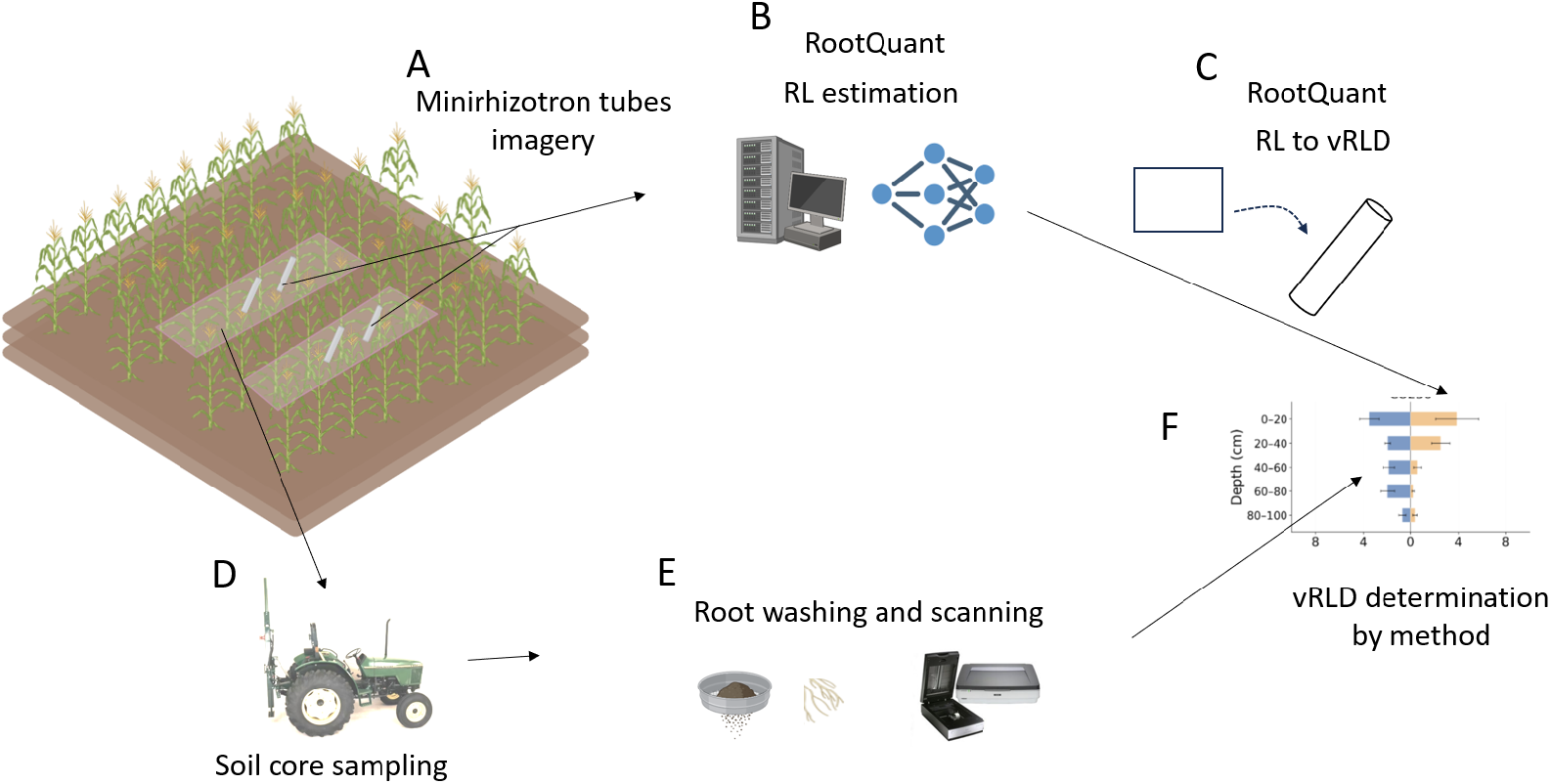
Overview of the field data collection, RootQuant inference pipeline, and field-scale vRLD validation workflow. (A) Minirhizotron images were acquired in field-installed transparent acrylic tubes using a Bartz BTC-100X camera system. (B) Individual RGB minirhizotron images were processed by the RootQuant deep learning framework to predict RL and RSA directly from raw imagery. (C) Image-level RL predictions were converted into vRLD estimates using the known imaging volume associated with each minirhizotron frame. (D) Independent soil-core samples were collected across field plots and soil-depth intervals for validation purposes. (E) Roots extracted from soil cores were washed, separated from soil material, and scanned to quantify root length using conventional root phenotyping procedures. (F) Minirhizotron image positions along obliquely installed tubes were converted into vertical soil depths, allowing RootQuant-derived vRLD estimates to be aggregated by depth strata and directly compared against soil-core-derived vRLD measurements across field plots and soil depths.

## 3 Results

The performance of RootQuant was evaluated at multiple levels, including image-level trait prediction, species-specific generalization, and transfer-learning ability.

Results show that the MS deep learning framework can jointly predict RL and RSA from minirhizotron imagery with high accuracy across multiple crop species without requiring pixel-level annotations or species-specific models (Table 1). For RL prediction, the model achieved length R^2^ = 0.90, RMSE = 2.86 mm, MAE = 1.08 mm, and for RSA, R^2^ = 0.88, RMSE = 4.16 mm^2^, MAE = 1.37 mm^2^ (Fig. 4).

**Table 1:** Performance of MS RootQuant for root length (RL) and root surface area (RSA) across three test subsets — all test images (MS), soybean-only, and maize-only — reported as R^2^, RMSE, and MAE. One erroneous soybean image (a WinRHIZO Tron labeling artifact with an anomalously large RSA residual) was excluded from the soybean-only RSA metrics; all RL metrics use the full test sets.

| Subset | Root length (RL) |  |  | Root surface area (RSA) |  |  |
| --- | --- | --- | --- | --- | --- | --- |
| | $R^2$ | RMSE (mm) | MAE (mm) | $R^2$ | RMSE (mm<sup>2</sup>) | MAE (mm<sup>2</sup>) |
| All test images (MS) | 0.90 | 2.86 | 1.08 | 0.88 | 4.16 | 1.37 |
| Soybean only | 0.88 | 3.46 | 1.32 | 0.87 | 4.85 | 1.64 |
| Maize only | 0.94 | 1.77 | 0.76 | 0.90 | 3.04 | 1.01 |

**Figure 4:**
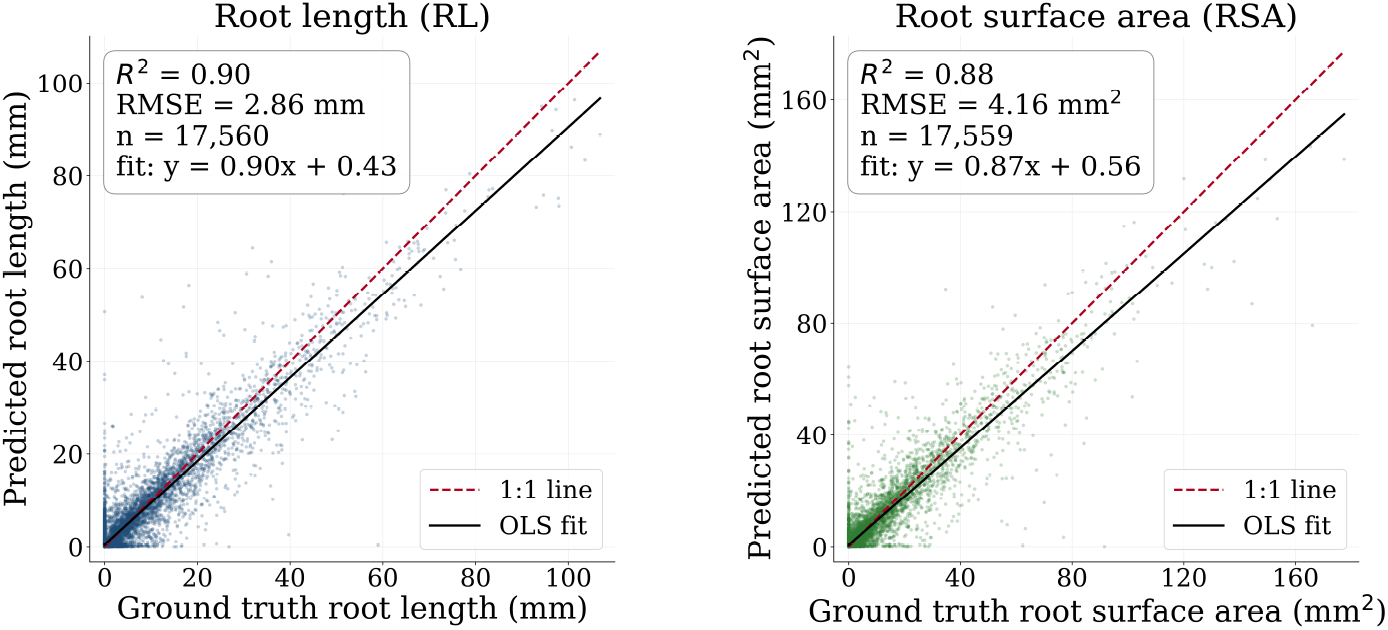
Predicted-vs-observed values of the MS RootQuant model for RL (left) and RSA (right) on the combined soybean and maize test dataset.

MS RootQuant achieved strong predictive performance across all test subsets (Table 1). For the combined dataset (MS, n = 17,560), RL yielded an R^2^ of 0.90, with an RMSE of 2.9 mm and an MAE of 1.1 mm. RSA showed lower accuracy (R^2^ = 0.88, RMSE = 4.2 mm^2^, MAE = 1.4 mm^2^), indicating that RL predictions were marginally more accurate and less variable than those for RSA. When evaluating the model’s predictions on individual crops, maize (n = 7,566) outperformed both the MS reference and soybean across all metrics: for maize, RL R^2^ reached 0.94 (RMSE = 1.8 mm, MAE = 0.8 mm) and RSA R^2^ reached 0.90 (RMSE = 3.0 mm^2^, MAE = 1.0 mm^2^). In contrast, performance on the soybean test dataset (n = 9,994) for RL was R^2^ = 0.88, RMSE = 3.5 mm, and RSA, R^2^ = 0.87, RMSE = 4.8 mm^2^. These results suggest that the mixed species MS dataset provides a balanced evaluation, whereas single crop subsets reveal distinct biases that affect prediction accuracy, particularly for soybean, despite its larger sample size.

The mixed species (MS) RootQuant model, trained on combined soybean and maize data, served as the baseline for cross species generalization (Fig. 5). On the maize test set, MS achieved strong performance for RL (R^2^ = 0.94, RMSE = 1.8 mm, MAE = 0.8 mm) and RSA (R^2^ = 0.90, RMSE = 3.0 mm^2^, MAE = 1.0 mm^2^). Compared to MS, zero-shot transfer (ZST) of the soybean-trained model on maize showed reduced performance across all metrics (RL R^2^ = 0.87, RMSE = 2.5 mm; RSA R^2^ = 0.80, RMSE = 4.2 mm^2^), highlighting the domain shift between species.

**Figure 5:**
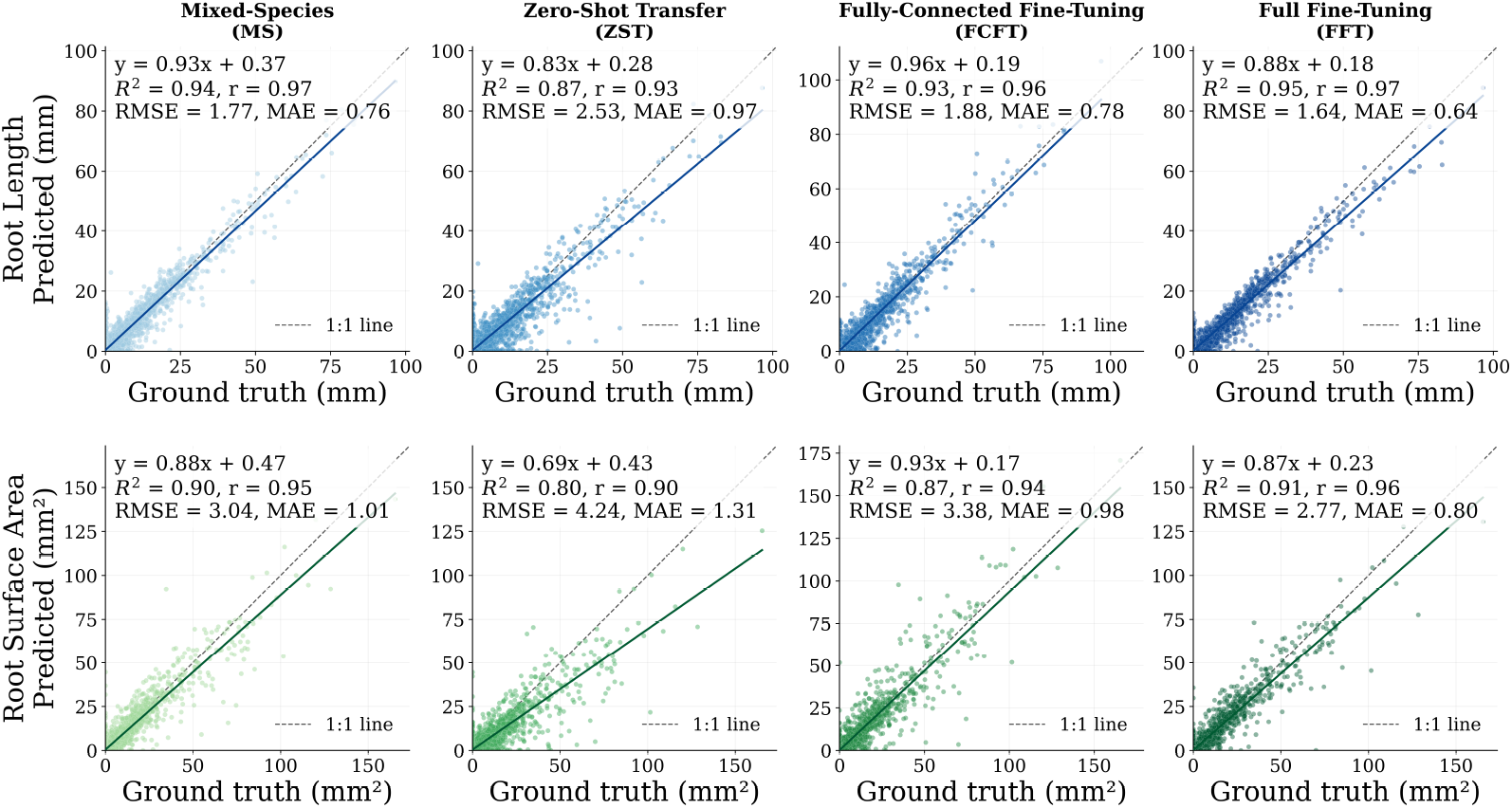
Predicted versus observed values for RL (top) and RSA (bottom) on the maize test set for (left to right) MS generalist, ZST, FCFT, and FFT. Each panel shows the OLS fit, R^2^, RMSE, and the 1:1 line.

Fine-tuning progressively closed this performance gap (Fig. 5,6). Fully connected-only fine-tuning (FCFT) improved performance relative to ZST (RL R^2^ = 0.93, RMSE = 1.9 mm; RSA R^2^ = 0.87, RMSE = 3.4 mm^2^), while full fine-tuning (FFT) yielded the highest accuracy for both RL (R^2^ = 0.95, RMSE = 1.6 mm) and RSA (R^2^ = 0.91, RMSE = 2.8 mm^2^). Notably, the MS model, despite being a single-weight generalist, remained competitive between FCFT and FFT, demonstrating its effectiveness for MS RootQuant applications without species-specific fine-tuning.

**Figure 6:**
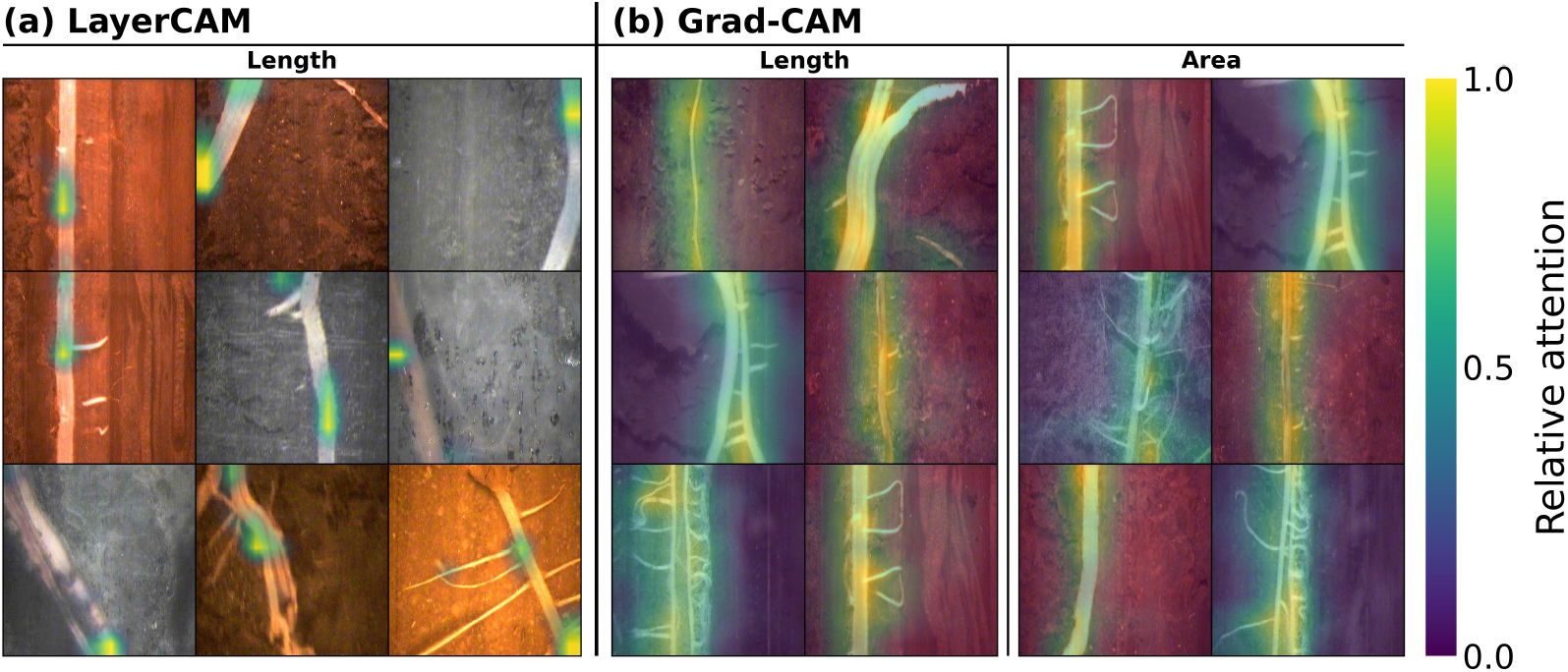
Quantitative LayerCAM maps (a) for RL and (b) Grad-CAM visualizations for (left) RL and (right) RSA predictions. Lower (blue) and higher (yellow) activation.

Quantitative attribution analyses demonstrated that RootQuant consistently focused on biologically relevant image regions. Across 726 images, LayerCAM attribution maps exhibited substantial enrichment within annotated root regions relative to area-matched random expectations (Fig. 6A), with approximately 2.6-fold and 2.9-fold increases in attribution energy for RL and RSA predictions, respectively. The higher mean enrichment observed on a per-image basis reflected the typical small fraction of image area occupied by roots.

Importantly, the localization signal remained robust under increasingly stringent spatial controls. Although the magnitude of enrichment decreased when length-preserving and area-preserving perturbations were introduced, attribution remained consistently concentrated within root structures, indicating that the observed signal could not be explained solely by spatial biases or image composition.

Complementary pointing-game analysis yielded a moderate absolute score (0.48), but this performance should be interpreted in the context of the sparse spatial occupancy of roots in an image. Relative to chance expectations, the probability that the highest-activation pixel coincided with a root region was approximately 2.5-fold greater than random placement, which is in agreement with the enrichment observed from attribution-energy analyses. The moderate pointing accuracy likely reflects the inherently diffuse spatial resolution of convolutional attribution maps rather than inaccurate localization.

Qualitative Grad-CAM visualizations further supported these findings, revealing broad activation patterns that consistently overlapped visible root structures (Fig. 6B). Together, qualitative and quantitative analyses indicate that RootQuant relies primarily on meaningful root features rather than background soil patterns or imaging artifacts, providing evidence that the model has learned relevant representations associated with root morphology.

### 3.1 Field-scale vRLD validation

RootQuant-derived vRLD was compared against vRLD derived from scans of roots washed out of destructively sampled soil cores across 15 field plots at each of 5 soil depths. vRLD was highest near the soil surface and declined with increasing depth (Fig. 7).

**Figure 7:**
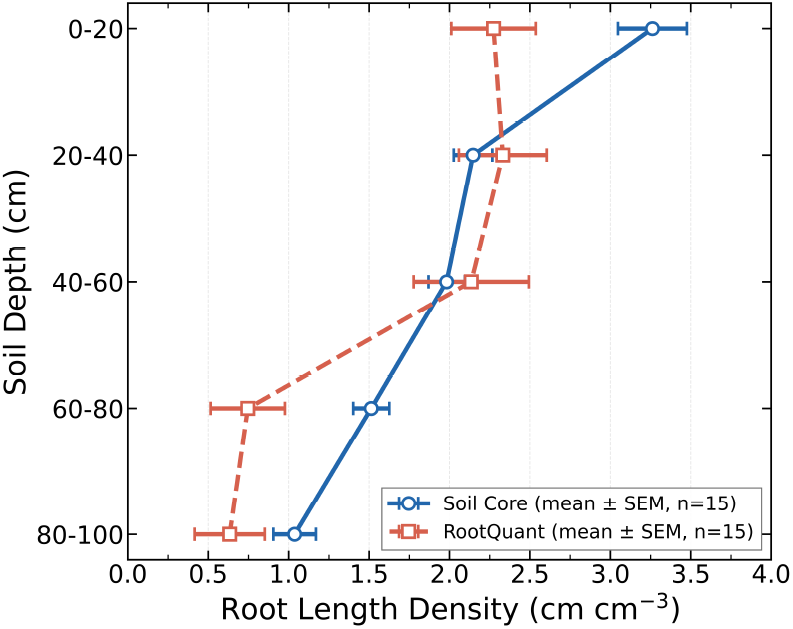
Comparison of vRLD between soil-core measurements and RootQuant predictions across five depth strata (0–100 cm, 20 cm intervals).

## 4 Discussion

Here, we introduce an end-to-end deep learning framework that simultaneously predicts RL and RSA directly from minirhizotron images using only whole-image numerical trait values as supervision. To our knowledge, this is the first multi-output model capable of jointly estimating these two complementary root traits without requiring per-pixel or skeleton-level annotations. By training on one of the largest reported maize and soybean minirhizotron datasets (89,186 human annotated images for training), RootQuant demonstrates that direct trait prediction from raw imagery is not only feasible but achieves high accuracy (RL R^2^ = 0.90, RSA R^2^ = 0.88 for the MS generalist model). For comparison, prior CNN-based studies were trained on far smaller hand-annotated sets — 50 images for the U-Net root segmenter of Smith et al. [20], 65 for SegRoot [21], 182 for faRIA [25], and roughly 1,500 for the automated pipeline of Bauer et al. [23] — placing RootQuant’s training set one to nearly three orders of magnitude above prior efforts. More importantly, our results reveal that a single MS generalist model can perform competitively across two major crop species, eliminating the need for species-specific model architectures or fine-tuning.

The results demonstrate that direct regression of RL and RSA from minirhizotron images is a viable alternative to conventional segmentation-and-skeletonization workflows. The MS generalist RootQuant model achieved strong predictive performance (RL RMSE = 2.9 mm, MAE = 1.1 mm; RSA RMSE = 4.2 mm^2^, MAE = 1.4 mm^2^) despite having no access to pixel-level root masks or skeleton annotations during training. This finding is particularly significant because most existing deep learning approaches for root phenotyping remain anchored in segmentation paradigms [19], [20], [21], [22], [23], [24] which require extensive manual annotation that is often unavailable for large historical datasets [23]. The practical advantage of our approach becomes clear when considering the nature of available minirhizotron archives. As noted earlier, decades of minirhizotron data stored in WinRHIZO Tron’s proprietary format contain only aggregated trait values per image, while the underlying pixel-level annotations are inaccessible. RootQuant circumvents this limitation entirely by treating the numerical trait values themselves as supervision signals. This design choice enables researchers to leverage existing historical datasets without re-annotation, dramatically expanding the pool of training data available for model development in the near future. In our case, access to 118,191 images with associated RL and RSA values—collected over an extended timeframe (i.e., 11 years) across the two major crop species—would have been impractical to re-annotate at the pixel level, yet these data proved sufficient to train a high-performing regression model.

However, the end-to-end proposed approach is not without limitations. The lower performance for RSA (R^2^ = 0.88) compared to RL (R^2^ = 0.90) suggests that surface area estimation may be more challenging to learn from whole-image supervision alone. RSA depends not only on root length but also on root diameter, which may be more variable across images and more susceptible to annotation noise in the original WinRHIZO Tron exports. Manual tracing operators may have been more consistent in marking root presence (affecting RL) than in precisely capturing diameter variations (affecting RSA). The Huber loss used during training was specifically chosen to mitigate the impact of such label noise, but residual inconsistencies remain. Future work could explore whether incorporating auxiliary self-supervised objectives [41] or multi-task learning with auxiliary losses (e.g., contrastive learning for root texture representation) [42] could improve RSA estimation without requiring pixel-level annotations.

The generalization evaluation revealed several important insights about the transferability of root trait prediction models across two relevant crop species. First, the MS generalist model—trained jointly on a large dataset of soybean and maize minirhizotron imagery—performed remarkably well on both species individually, achieving an RL R^2^ of 0.94 on maize and 0.88 on soybean with a single set of weights. This performance exceeded the zero-shot transfer of a model trained only on soybean to maize (ZST: RL R^2^ = 0.87, RSA R^2^ = 0.80), demonstrating that MS training confers genuine generalization benefits rather than only single species-specific features.

The zero-shot transfer results account for the domain shift between soybean and maize minirhizotron imagery. The performance drop from RL R^2^ = 0.94 in MS to RL R^2^ = 0.87 in ZST on the maize test set represents a relative decrease of approximately 7.4%. This result indicates that a model trained exclusively on soybean does not fully generalize to maize, despite the two species being imaged by the same research team using consistent protocols at field sites in close proximity with common soil types. This domain shift likely arises from differences in root morphology (maize produces thicker, more structurally distinct roots compared to soybean’s finer root system) [5], [43] and possibly phenological timing of image acquisition [7].

Encouragingly, both fine-tuning strategies successfully reduced the domain shift gap. FCFT, which updated only the regression head while keeping the soybean-trained backbone frozen, improved RL R^2^ from 0.87 to 0.93—achieving performance nearly equivalent to the MS model (R^2^ = 0.94). This finding suggests that the backbone convolutional features learned from soybean images are largely transferable to maize, and that species-specific differences can be accommodated primarily through recalibration of the final regression layers. FFT provided a marginal additional improvement (RL R^2^ = 0.95), but the small gain relative to FCFT (0.93 to 0.95) may not justify the additional computational cost and risk of catastrophic forgetting in practical applications.

From a practical standpoint, these results indicate that a MS training strategy is the most efficient path to a generalist model that performs well across species without fine-tuning – at least when the species have similar fine root structures and are growing in common soil conditions. For researchers working with a new crop species not represented in the training data, the FCFT strategy offers a computationally efficient adaptation method that requires only a modest set of labeled target-species images to retrain the regression head. The fact that FCFT nearly closed the domain gap suggests that root feature representations are surprisingly transferable across species, at least between maize and soybean. We are optimistic this strategy may transfer to other crops and herbaceous species in locations with strong seasonality. However, the likelihood that it breaks down will be greater if attempting to work in woody species and/or aseasonal environments where fine root demographic patterns may be very different [44].

An important consideration for end-to-end DL regression models trained with weak whole-image, single-trait-value supervision is their susceptibility to shortcut learning, whereby predictive performance may arise from spurious correlations in the background rather than from biologically meaningful structures [38]. Our analysis provides evidence that this is not the dominant mechanism underlying RootQuant predictions. Quantitative LayerCAM analysis showed that attribution was significantly higher within annotated root regions than in multiple null models. Qualitative Grad-CAM visualizations further support these findings, with the most influential regions for prediction consistently overlapping visible roots rather than surrounding soil or imaging artifacts. Overall, these results indicate that RootQuant learns representations predominantly grounded in meaningful root structures despite the absence of explicit pixel-level supervision. Nevertheless, the attribution evaluation provides only indirect evidence of the model’s internal representation and cannot completely exclude residual contributions from non-root image features.

Relative to Khoroshevsky et al. [26], [27], who demonstrated direct root length prediction from minirhizotron images in grapevine, our work extends the end-to-end regression paradigm. First, we jointly predict both RL and RSA, recognizing that these traits capture complementary aspects of root characteristics. Second, we train and evaluate on a substantially larger dataset — 118,191 images versus 531 in the grapevine study of [26] and 4,015 in the multi-crop study of [27] — spanning two major crop species and multiple years, providing stronger evidence for generalization. Compared to segmentation-based deep learning approaches [19], [20], [21], [22], [23], [24], [25], RootQuant trades pixel-level accuracy for scalability and historical data compatibility. Moreover, such models require manual segmentation masks for training, which are unavailable for most historical minirhizotron datasets. For researchers sitting on years of WinRHIZO Tron exports, RootQuant offers a path to automate trait extraction without re-annotation. The two approaches are not mutually exclusive: future work could use RootQuant predictions to pre-screen images for variation in RL and RSA, and then perform targeted segmentation on a subset of images for detailed semantic segmentation analysis.

It has long been recognized that different methods of assessing fine root system size and depth distribution – including minirhizotrons, soil coring and trenching – have different strengths and weaknesses, but can deliver generally consistent results even if the inherent noise in all data types prevents very strong correlations among methods [45], [46], [47], [48]. Minirhizotrons are especially favored when quantifying temporal time courses of fine root length and distribution is a research goal, and this is made much more feasible when automated image analysis is available [47], [48], [10]. However, minirhizotron analysis can fail to reproduce the vertical profile of vRLD when the quality of minirhizotron access tube installation is poor [7], [46]. Therefore, we confirmed that both RootQuant-derived vRLD and soil-core measurements captured the expected vertical gradient in root density, with highest values in the 0–20 cm surface stratum decreasing progressively to the 80–100 cm depth. Larger scale comparison of methods across more diverse test plots should be performed in the future to provide a more formal quantitative evaluation than the modest test that was possible in the current study. Several limitations of this study should be acknowledged. First, the training labels (RL and RSA) originated from manual tracing in WinRHIZO Tron, which itself has known limitations including operator subjectivity, difficulty resolving overlapping roots, and systematic underestimation of fine roots [7], [10], [49]. RootQuant inherits these biases; any systematic errors in the original WinRHIZO Tron labels are learned as ground truth. Second, the dataset included only maize and soybean; generalization to other species (e.g., wheat and perennial crops) with different root characteristics remains untested. Third, the imaging volume for vRLD conversion assumed a uniform depth-of-field of 2 mm, but the actual soil volume visualized may vary with soil moisture, compaction, or root proximity to the tube [7].

Future work should explore several avenues for improving model performance and broadening the applicability of RootQuant. Incorporating unlabeled or weakly labeled data through self-supervised pretraining [41], [50] could improve feature learning. Domain adaptation methods beyond fine-tuning, such as unsupervised adversarial alignment [51], could reduce the need for labeled target-species data. Additionally, extending RootQuant to predict additional traits (e.g., mean root diameter, root volume, or root color as a proxy for age) would increase its utility for root studies.

## 5 Conclusion

RootQuant demonstrates that end-to-end prediction of root length and surface area from minirhizotron images is feasible, accurate, and scalable using only whole-image numerical trait values as supervision. Trained on 118,191 images spanning maize and soybean across several seasons, the MS generalist RootQuant model achieves RL R^2^ = 0.90 and RSA R^2^ = 0.88 on held-out test data, with strong cross-species generalization requiring no fine-tuning for maize. The model’s learning process focuses on meaningful root biological features. The model’s RL predictions convert meaningfully to vRLD, showing qualitative and quantitative agreement with independent soil-core measurements across soil depths, particularly in the 20–60 cm interval. By eliminating the need for pixel-level annotations, RootQuant unlocks decades of historical minirhizotron data previously inaccessible to deep learning approaches, enabling large-scale retrospective analyses and accelerating field-based root phenotyping for crop improvement programs.

## Acknowledgments

We thank all members of the Leakey Laboratory for their valuable contributions and dedication to the implementation of field trials, data collection, and other essential activities. In particular, we express our deep gratitude for their effort in the manual tracing of roots across this extensive dataset between 2009 and 2020; without such a team effort, this work would not have been possible.

